# Adaptive stretching of representations across brain regions and deep learning model layers

**DOI:** 10.1101/2023.12.01.569615

**Authors:** Xin-Ya Zhang, Sebastian Bobadilla-Suarez, Xiaoliang Luo, Marilena Lemonari, Scott L. Brincat, Markus Siegel, Earl K. Miller, Bradley C. Love

## Abstract

Prefrontal cortex (PFC) is known to modulate the visual system to favor goal-relevant information by accentuating task-relevant stimulus dimensions. Does the brain broadly re-configures itself to optimize performance by stretching visual representations along task-relevant dimensions? We considered a task that required monkeys to selectively attend on a trial-by-trial basis to one of two dimensions (color or motion direction) to make a decision. Except for V4 (color bound) and MT (motion bound), the brain radically re-configured itself to stretch representations along task-relevant dimensions in lateral PFC, frontal eye fields (FEF), lateral intraparietal cortex (LIP), and inferotemporal cortex (IT). Spike timing was crucial to this code. A deep learning model was trained on the same visual input and rewards as the monkeys. Despite lacking an explicit selective attention or other control mechanism, the model displayed task-relevant stretching as a consequence of error minimization, indicating that stretching is an adaptive strategy.

---

Adaptive agents, whether biological or artificial, configure themselves for the learning and decision task at hand. For example, when searching for one’s car keys, one might focus on features consistent with the shape and metallic sheen of a key. Prefrontal cortex (PFC) modulates the visual system to favour goal-relevant information [1, 2]. Goal-directed attention can reconfigure the visual system to highlight task-relevant features and suppress irrelevant features [3]. Over longer time horizons, learning processes build internal representations that reflect these task pressures [4, 5] with vmPFC critical for determining which aspects of the current context are relevant [6]. Attention can be viewed as *stretching* representations along relevant dimensions [7, 8, 9], which is reflected both in behavior and brain response [5, 10, 11]. Stretching accentuates differences along goal-relevant dimensions while minimizing differences along goal-irrelevant dimensions. For example, when searching for one’s car keys, objects varying in metallic sheen should become more dissimilar.

One possibility is that the brain radically re-configures itself across regions to optimize for the current task. Consistent with this possibility, effects of endogenous attention have been observed across the visual cortical hierarchy [12, 13, 14], including as early as V1 [15, 16]. Alternatively, some areas devoted to modality specific processing, such as the middle temporal cortex (MT) for movement direction and visual area V4 for object properties like color, may be invariant across task contexts.

The ideal study to evaluate whether radical reconfiguration occurs would record from multiple brain sites while cueing the relevant dimension for a categorization decision on a trial-by-trial basis. We analyzed spiking data from a study[17] that met these criteria. On each trial, the rhesus monkeys viewed a cue that indicated whether color or direction of movement was the relevant dimension for the decision (Fig. 1a). Once cued, colored dots moved with 100% coherency in a given direction. The monkeys responded by moving their eyes left or right, depending on the value of relevant (either color or motion) stimulus dimension (Fig. 1c). Recording sites included lateral prefrontal cortex (PFC), frontal eye fields (FEF), lateral intraparietal cortex (LIP), inferotemporal cortex (IT), V4, and MT. Unlike previous fMRI studies [5, 11] that considered stretching along relevant dimensions during categorization decisions, monkey multi-unit spiking data affords the possibility of evaluating whether spike timing, over and above spiking rate, is a critical component of the neural code.

**Fig. 1.**
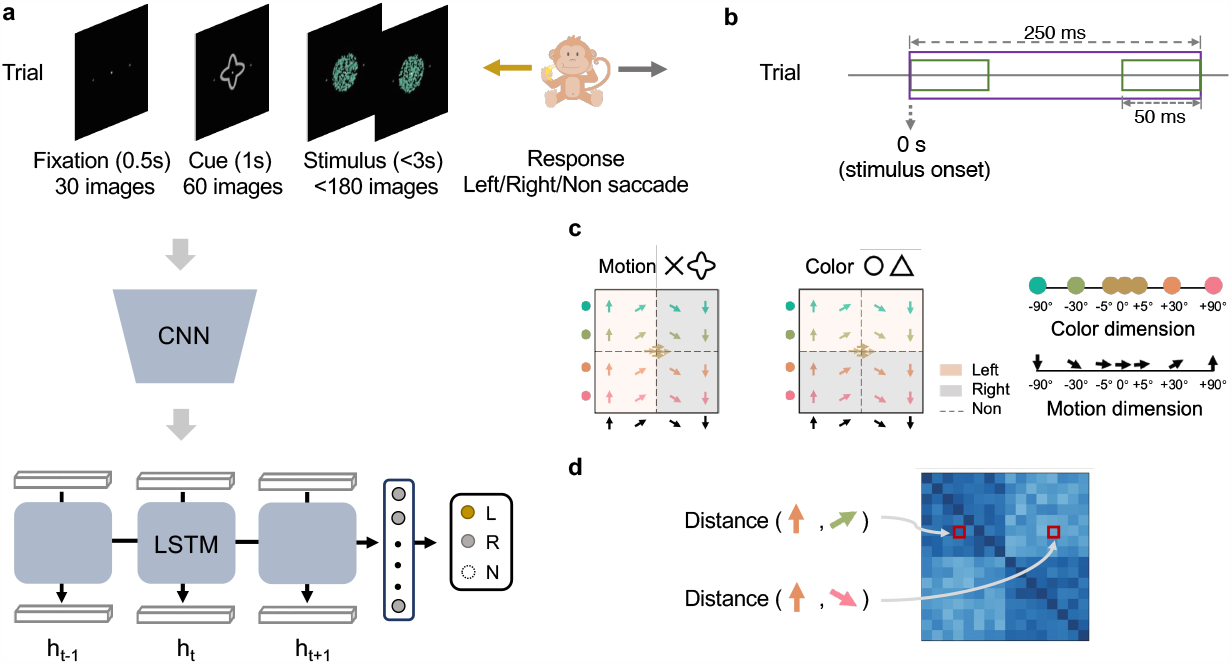
Overview of the behavioral task and CNN-LSTM modeling. **a**, Monkeys were shown images depicting the fixation point (0.5 s), cue (1 s), and stimulus (up to 3 s). During each trial, monkeys held central fixation until they responded by making either a left or right saccade. They were not to respond on ambiguous trials. These same images were shown to the ANN model consisting of a pretrained CNN for visual processing and a stacked (i.e., multilayer) LSTM that learned how to appropriately respond given the task cue and stimulus. Like the monkeys, the CNN-LSTM learned through trial-and-error to respond left (L), right (R), or to withhold (N) a response on ambiguous trials. **b**, Spiking data were analyzed over a 250 ms period (shown in purple) that began at stimulus onset. Analyses either considered this time period as a whole or in 50 ms overlapping segments using a sliding window (shown in green) that was 50 ms wide and moved in 10 ms steps. **c** There were 21 color-motion stimuli constructed from the color and motion space. Five stimuli were ambiguous at the origin of the color-motion space. The remaining 16 stimuli were evenly spread across the four quadrants. A cross or quatrefoil cue indicated that motion was relevant to the decision. A circle or triangle cue indicated the color was relevant. In both task contexts, the monkeys responded with a leftward or rightward saccade. **d**, From the 16 non-ambiguous trials, we constructed 16x16 dissimilarity matrices (i.e., RDMs) for data analyses. To determine the value of an entry in the RDM, the spiking measures for two items would be compared using a particular distance function (e.g., ISI, Euclidean distance, etc.) and the value was stored in the row and column corresponding to the two items. Likewise, RDMs were made for the ANN using the activity patterns from LSTM’s hidden unit activations.

Considering goal-relevant stretching along a relevant dimension in non-human animals offers opportunities to examine how control systems themselves can be learned and configured absent language instruction. Non-human animals, like artificial neural networks (ANNs), are naive to these laboratory tasks and, unlike verbally instructed humans, have to learn how to allocate their attention through trial-and-error.

Nevertheless, non-human animals, including rats, can learn to selectively attend to relevant dimensions in related paradigms [18].

While models of goal-directed attention contain dedicated control systems that selectively weight relevant dimensions [7, 9, 8, 19, 20], one possibility is that these control systems themselves can be learned by non-human animals and ANNs, enabling them to reconfigure themselves in response to task cues. According to this hypothesis, brains and ANNs are powerful statistical learning machines that build control structures that adaptively stretch learnt representations along relevant dimensions to facilitate task performance. In effect, these systems may build the cognitive machinery that is presupposed in previous work.

To evaluate this possibility, we constructed a ANN consisting of a deep convolutional neural network (CNN) and a stacked long short-term memory (LSTM). The CNN was pretrained to perform object recognition on naturalistic images [21]. The CNN part of the ANN, akin to the monkeys’ visual system [22, 23, 24, 25, 26], was assumed to be developed and stable prior to the study. Thus, its weights were fixed. The sequences of images forming a trial (e.g., the task cue and the moving dots; Fig. 1a) were fed into the CNN, whose outputs served as inputs to the LSTM [27]. The LSTM is a recurrent network that can process time series to make the left/right decision the monkeys did (see Supplementary Information for details). Like the monkeys, the LSTM part of ANN learned through trial-and-error from sequences of images, absent verbal instruction.

Of course, the brain embodies constraints that ANNs do not. Brain regions may be limited to processing certain stimulus dimensions and may lack the flexibility to reconfigure. For example, in the present investigation, V4 and MT may remain bounded to processing color and motion, respectively. However, to the extent the brain can be viewed as an overparameterized statistical learning engine [28], the ANN and brain should converge. We predict this convergence will be realized by representations stretching along the goal-relevant stimulus dimension. Observing stretching in the ANN would provide a formal account of how control and top-down attentional mechanisms can arise from simply maximizing task performance, as opposed to relying on preordained mechanisms.

To foreshadow the results, we observed stretching across all layers of the ANN’s LSTM and in the brain, except for areas V4 and MT, which were modality bound. We also used a cognitive model with a dedicated attention mechanism to assess stretching in the brain data, which corroborated these conclusions. Finally, we analyzed the similarity and fidelity of neural representations [29] and discovered that spike timing was critical to how the brain coded representations of the stimuli and task.

## Results

### Spike timing and neural representation

One initial question is which measure of neural similarity is most aligned with the neural recordings [29]. To answer this question, we assessed various measures of neural similarity to determine which one maximized representational similarity [30] with the experimenter-defined stimulus coordinates of color and motion. The high temporal resolution of the spike trains allowed us to consider whether dissimilarity measures that take into account spike timing, such as ISI [31] and SPIKE [32], offer advantages over rate-based measures, including Euclidean, cosine, and Pearson measures. ISI and SPIKE did surpass the non-timing measures (Fig. 2). To confirm the significance of the type of measure, we conducted a one-way ANOVA analysis encompassing all measures, *F* (4, 105) = 143.06, *p <* .0001. In particular, ISI performed best and will be used throughout the remaining analyses in this contribution (see Bonferroni corrected *t* -tests in Extended data Tables 1, 2). ISI emphasizes the relative intervals between spikes, whereas SPIKE also incorporates the absolute timing of spikes which can be useful for evaluating synchrony between spike trains (see Methods and Supplementary Information). The advantage of spike timing measures over rate-based coding held across all recording sites (Extended Data Figs. 3 and 4).

**Fig. 2.**
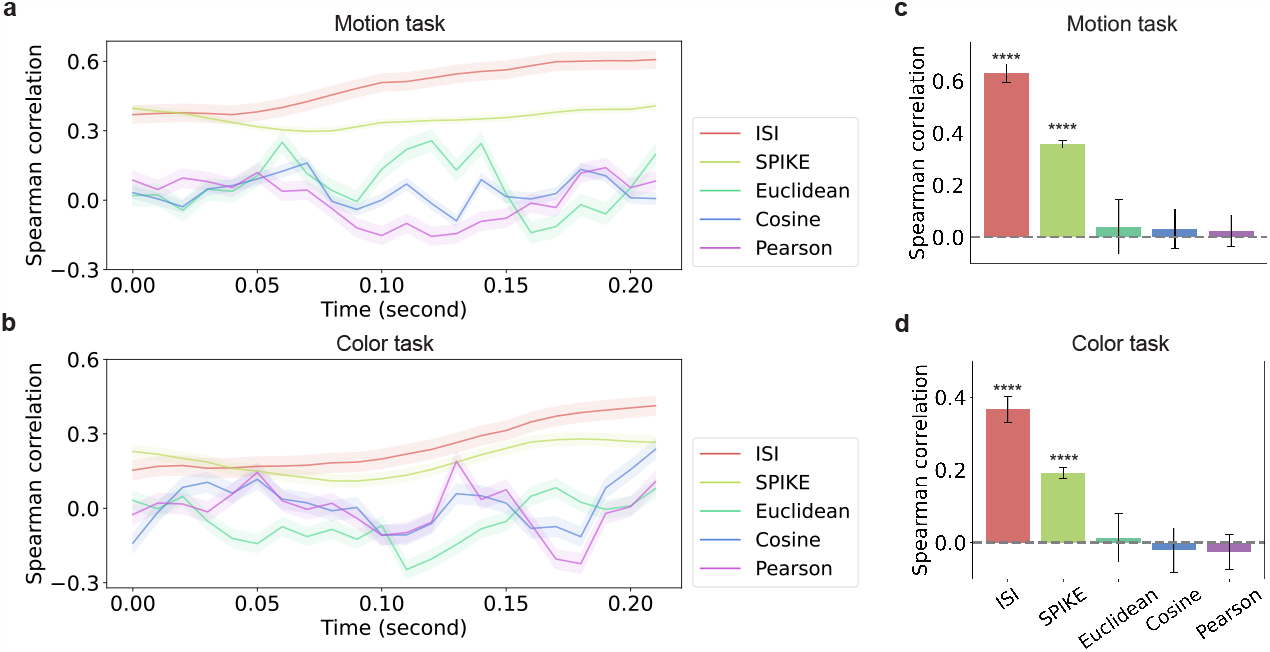
Spike timing measures best capture the experimenter intended coordinates. We compared the Spearman rank correlation between dissimilarity matrices (i.e., RDMs) constructed from the monkey data for the 16 unambiguous items (see Fig 1d) with an RDM derived from the experimenter-intended stimulus coordinates (see Fig 1c). Higher correlation indicates better agreement. **a**, When motion was relevant, the timing measures ISI and SPIKE surpass rate coding measures and best capture the experimenter-intended coordinates. **b**, The same pattern was found when color was relevant to the monkey’s decision with ISI once again proving best. The band around each dissimilarity measure depicts the 95% confidence intervals. Time on the horizontal axis is measured from stimulus onset (see Fig 1b). **c, d**, The same pattern of results holds when the entire 250 ms time period is analyzed as a whole.

### Dimensional stretching found in both brain and model activity

One key question is whether the brain radically re-configures itself across regions to optimize for the current task, which in the present study would manifest as stretching along the relevant dimension (color or motion) on each trial. Likewise, we consider whether the LSTM, simply by maximizing performance absent an explicit control or attentional mechanism, will adaptively stretch its representations.

Stretching was assessed by considering item pairs that mismatched on one dimension and matched on the other. For example, we predict items mismatching on color and matching on motion should be more dissimilar when color is relevant than when motion is relevant. The dissimilarity of the 24 (*i, j*) item pairs that mismatch on color and match on motion is denoted *D*_*c*_(*i, j*)^*c*^ when color is relevant and *D*_*c*_(*i, j*)^*m*^ when motion is relevant. The 24 item pairs that mismatch on motion and match on color is denoted *D*_*m*_(*i, j*)^*c*^ and *D*_*m*_(*i, j*)^*m*^ for when color and motion are relevant, respectively. Stretching should be greatest when items mismatch on the relevant dimension.

This prediction was confirmed both in the brain and LSTM activity (Fig. 3). Mismatches along a stimulus dimension were more consequential when that dimension was task relevant. In the monkey data, *D*_*c*_(*i, j*)^*c*^ − *D*_*c*_(*i, j*)^*m*^ and *D*_*m*_(*i, j*)^*m*^ − *D*_*m*_(*i, j*)^*c*^ were significantly different than 0, *M* = 0.0007, *t*(23) = −4.274, *p <* .0001 and *M* = 0.0007, *t*(23) = −4.132, *p* = 1.4*e* − 4, respectively. In the LSTM simulations, *D*_*c*_(*i, j*)^*c*^ − *D*_*c*_(*i, j*)^*m*^ and *D*_*m*_(*i, j*)^*m*^ − *D*_*m*_(*i, j*)^*c*^ were significant different than 0, *M* = 4.738, *t*(23) = −7.259, *p <* .0001 and *M* = 7.0694, *t*(23) = −9.887, *p <* .0001), respectively.

**Fig. 3.**
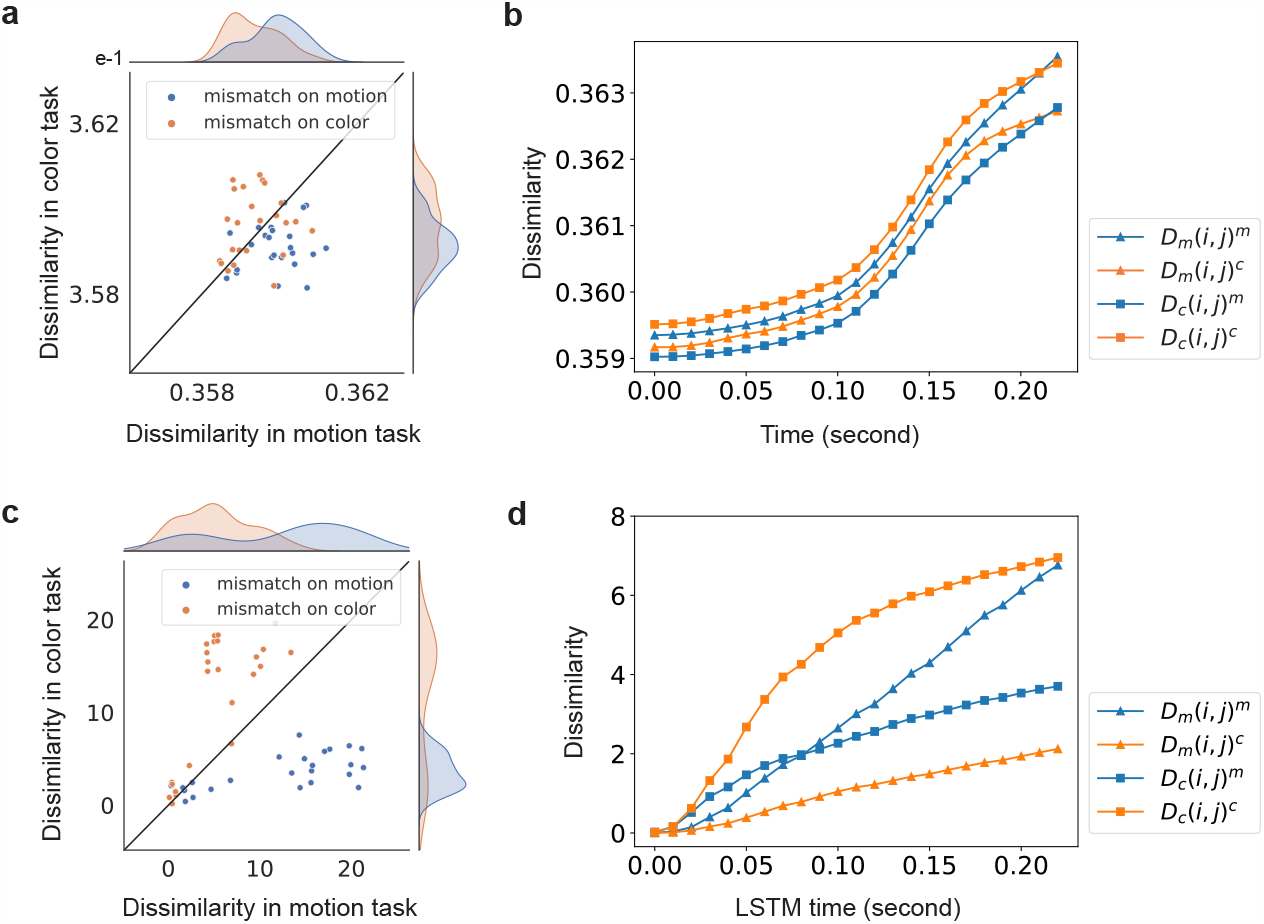
Dimensional stretching occurs in both neural data and model representations. **a**, Dissimilarity between item pairs mismatching on motion or color is increased when that dimension is task relevant. The density distributions of these dissimilarities also indicate this task-modulation. **b**, Changes in dissimilarity over trial time (time 0 s is stimulus onset). For stimulus pair (*i, j*), the underscore index indicates the dimension they mismatch, e.g., *D*_*c*_(*i, j*) indicates items *i* and *j* mismatch on color and match on motion. Item pairs that mismatch on motion are more different in the motion-relevant context than in the color-relevant context. Mirroring, item pairs that mismatch on color are more different in the color-relevant context than in the motion-relevant context. **c, d** The CNN-LSTM model shows the same qualitative pattern of performance – stimulus pairs that mismatch on the task relevant dimension were most dissimilar.

### Stretching across model layers and brain regions

To assess how the six brain regions and LSTM model layers are modulated by task, we applied a cognitive model adapted from the generalized context model [8]. The model fit assessed how much attention, *w*_*r*_ (between 0 and 1), is dedicated to the task-relevant dimension with the irrelevant dimension weighted by 1 − *w*_*r*_ (see Methods). This cognitive model assumes the two-dimensional psychological space shown in Figure 1c and is useful for estimating the best fitting attention parameter *w*_*r*_, which characterizes representational stretching.

Figure 4a shows how brain regions representation spaces stretch along the relevant dimension, except for MT and V4. To assess these effects, we conducted permutation tests in which dimension considered relevant was shuffled (1000 times to construct the null distribution). For the PFC, FEF, LIP and IT, *w*_*r*_ was significantly larger than expected in both tasks contexts, all p-values less than .01. However, MT is modality bounded and focuses on motion dimension in both task contexts, whereas V4 focuses solely on the color dimension. Accordingly, these two modality bound regions had higher *w*_*r*_ values for their preferred dimension and lower *w*_*r*_ for their non-preferred dimension as compared to the null distribution, all p-values less than .01. For the LSTM, apart from earliest layer (*p* = .33), all layers had *w*_*r*_ values greater than expected by chance, all p-values less than .05 (see Extended Data Tables 3 and 4).

**Fig. 4.**
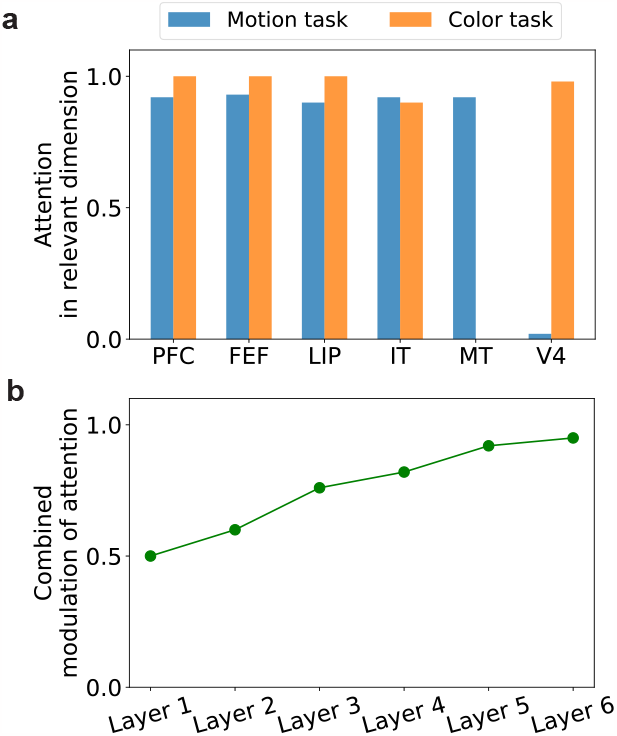
Task-relevant attention allocation in brain regions and LSTMs layers. **a**, The allocation of attention (estimated by fitting a cognitive model) across the six brain regions is shown. MT prioritizes motion and V4 color regardless of the task context. **b**, Fitting the cognitive model to LSTM representations revealed that more advanced layers devoted more attention to the relevant stimulus dimension.

### LSTM re-configures itself based on the task goal unlike MT and V4

There is no straightforward layer-to-brain correspondence between LSTM layers and brain regions (Fig. 5). Whereas MT and V4 appear dedicated to processing motion and color, respectively, LSTM layers freely reconfigure as a function of the task context. For example, V4 shows a decent correspondence with LSTM layers when color is relevant, whereas MT does when motion is relevant. Unlike some brain regions, the LSTM is more flexible and can fully reconfigure itself depending on the task.

**Fig. 5.**
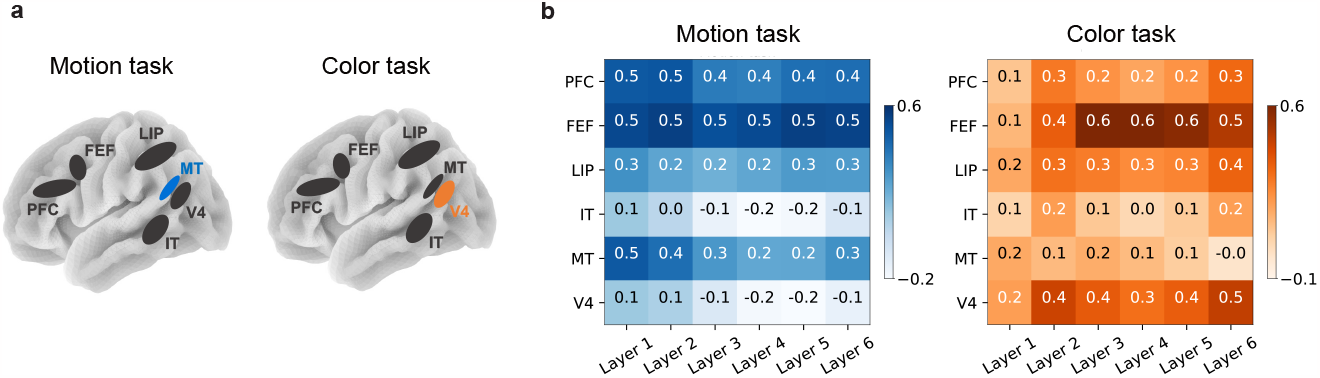
Alignment between brain region and LSTM layers. **a**, One hypothesis, supported by our previous analyses, is that recording sites, with the exception of MT (motion fixated) and V4 (color fixated), should align equally well with LSTM activity in the motion and color tasks. **b**, Alignment, as measured by the Spearman correlation of the RDMs (Fig. 1d) for brain and model activity, is shown. Most brain regions generally aligned across model layers and tasks. IT was generally misaligned. In contrast, MT better aligned in the motion task, whereas V4 better aligned in the color task. Consistent with previous analyses, MT and V4 do not seem to reconfigure as a function of task.

## Discussion

We considered whether the brain re-configures itself to optimize for the current task. For example, do neural representations of stimuli stretch along dimensions relevant to the task? We analyzed data from a task that, depending on a visual cue at the beginning of each trial, required monkeys to selectively attend to one of two stimulus dimensions to make a decision. We found support for the hypothesis that the brain radically re-configures itself to optimize performance by accentuating differences along task-relevant dimensions. An ANN that was trained on the same visual information as the monkeys also displayed task-relevant stretching as a consequence of trying to reduce error, indicating that stretching is an adaptive strategy that can arise in systems that aim to optimize performance.

Stretching was established in the monkey data by analyses that compared stimulus pairs matching or mismatching on a relevant dimension. Pairs that mismatched on a relevant dimension were more dissimilar than those that mismatched on an irrelevant dimension (Fig. 3). We also evaluated stretching by using a model-based approach in which a cognitive model with a selective attention parameter was fit to the neural data (Fig. 4). Stretching was found at all recording sites, except for V4 and MT, which were bound to processing color and direction information, respectively.

One way ANN activity diverged from the brain data is that the ANN’s LSTM layers freely reconfigured themselves as a function of the task context. In other words, unlike the brain, the ANN was not modality bound. Accordingly, there was no straightforward relationship between brain regions and LSTM layers. For example, when color was relevant, V4 and network activity correlated, whereas when motion was relevant, MT and network activity correlated (see Fig. 5).

Finally, although not a primary hypothesis, the monkey spiking data afforded the possibility of evaluating whether spike timing or rate-based measures better characterize the neural code. Spike timing did appear relevant to how the brain represented stimulus information (see Fig. 2). These findings suggest a roll for spike timing that should be further explored, including in computational models. While our ANN was a recurrent network that processed the sequences of images forming a trial through time, the activity of each of its units was a scalar akin to rate-based coding. New models will need to developed to assess hypothesis concerning the functional role of spike-timing information in decision making.

One strength of the modeling approach is that the ANN was applied to the same image sequences forming a trial as those the monkeys experienced. Rather than attempt to incorporate biological constraints, we treated the model as a general-purpose statistical machine and observed how it tailored itself to the task. Under this approach, convergences between the ANN and the monkeys may reflect general information processing principles, whereas divergences may reflect unique properties of biological systems.

We found that the ANN performed as if it had a cognitive control mechanism that reconfigured network layers based on the task cue, stretching representations along task-relevant dimensions. In contrast, cognitive models, like the model we used in our analyses (Fig. 4), have dedicated attentional mechanisms. The ANN results suggest that general-purpose learning systems can learn to control themselves absent purpose-built control circuits. In terms of divergences, we found that, unlike V4 and MT, the ANN was not modality bound by layer. This result suggests that there is a benefit or constraint in biological systems for localized processing of perceptual streams or dimensions.

Although this is just one study and model, we believe this modeling approach could complement other approaches in the literature, such as approaches that aim to test a priori hypotheses regarding correspondences between brain regions and model layers as is commonly done in the object recognition literature [22, 23, 24, 25, 26, 33]. Our approach may prove useful in other domains, including the meta-learning challenge of “learning to learn”. In particular, one interesting question is whether the implicit control structures the ANN developed for the present task would provide a useful starting point for mastering new tasks that involve selective attention. There might be a great deal to learn about the brain by considering the computational challenges it faces.

## Acknowledgements

This work was supported by ESRC (ES/W007347/1), Wellcome Trust Investigator Award WT106931MA, and Royal Society Wolfson Fellowship 183029 to B.C.L, JPB Foundation, Picower Institute for Learning and Memory, and Office of Naval Research N00014-23-1-2768 to EKM, and China Scholarship Council (Grant No. 202206260103) to X.Y.Z.

## Data and Code availability

The codes used in this work are available at https://github.com/xinyacheung/neural_similarity. All behavioral and electrophysiological data are archived at the Centre for Integrative Neuroscience, University of Tübingen, Germany.

## Methods

### Experiment and simulation description

In this work, we aim to model monkey brain learning on a flexible visuomotor decision-making task with our CNN-LSTM architecture. In the original experiment [17], two rhesus (one male and one female) monkeys were shown a series of stimuli videos and learned to categorize the color or the orientation of stimuli dots based on cues they saw. Recordings were performed using three stereotactically positioned recording chambers, which covered the frontal cortex (FEF and PFC), parietal cortex (LIP), and occipitotemporal cortex (IT, MT and V4). There were 21 color-motion stimuli (5 stimuli on or near the category boundary and 16 stimuli evenly spread across the four quadrants) from 7 possible colors and 7 possible motion directions (Fig. 1c). On each trial, monkeys saw one stimulus (dot pattern), that is, all dots with the same color moved 100% coherency in the same direction. Depending on the task cued at the beginning of each trial, the monkeys categorized either the color (red vs. green) or motion direction (up vs. down) of the stimulus and reported their percept with a left or right saccade. We did our best to replicate the training procedure as faithfully as possible by tasking our model with the same categorization problems, using identical fixation, cue and stimulus images (dot pattern) and training our model in a trial-by-trial fashion. We detail how experiments were performed while highlighting and justifying the necessary adjustments we have made in Supplementary Information.

To model the learning process and relate the neural data of trained monkeys, we employ training trials and test trials into our model and predict responses. We have a total of 6720 trials covering 4 cues, 2 speeds, 21 color-motion stimuli with 40 random stimulus dots’ initial positions (4 *×* 2 *×* 21 *×* 40 = 6720) and the training/test trial is 3 : 1 (i.e., 5040 training trials and 1680 test trials involved 30 and 10 random initial positions, respectively). Notice that although these trials use the same color-motion stimulus (dot pattern), the dots’ initial position in each trial is random, exactly as the original experiment. Matched with the original experiment, we include ambiguous trials of five boundary stimuli in our training and test but excluding them from all data analyses, that is we only consider 16 color-motion stimuli in our analyses. We use L (left) to represent the categorization of stimuli with upward motion in the motion task or stimuli made of greenish dots in the color task as the same class. Likewise, we use R (right) to represent the categorization of stimuli with downward motion in the motion task or stimuli made of reddish dots in the color task as the other class. We use N (no response) to represent ambiguous trials in training ANN like monkeys did.

### CNN-LSTM architecture

We tailor a CNN-LSTM framework containing a deep convolutional neural network (CNN) and a stacked long short-term memory neural network (multilayer LSTM). The overall learning task of our model is to categorize the same set of stimuli (a left or right saccade based on either motion or color and no response if ambiguous trials), which is identical to the training procedure as monkeys did. The CNN front-end is used to simulate the monkey visual system and the stacked LSTM is to simulate multiple brain regions such as MT, V4 which might be involved in the learning task.

Formally, we denote the CNN front-end as **f** which in our case is a pre-trained VGG-16 up to the pre-softmax layer. Given a trial of *n* stimuli, *T*_*i*_, **f** transforms raw images into representations *Q* ∈ ℝ^*n×d*^, where *d* is the size of the pre-softmax layer,

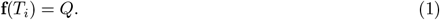

We then denote the stacked LSTM as **g** which transforms the visual representations *Q* into *H* ∈ ℝ^*d×h*^, following

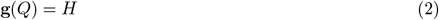

where *h* represents the number of LSTM cells in each layer. Finally, to transform LSTM representations into the decision response, we apply two linear transformations *W*_1_ ∈ ℝ^*h×ht*^ and *W*_2_ ∈ ℝ^*ht×m*^, mapping *H* to *P* ∈ ℝ^*n×m*^ where each row of *P* being a probability distribution over the *m* classes that the original experiment was trained to classify,

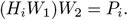

In our simulations, we employ 270 images in each trial (*n* = 270), extract 1000 high-level features from VGG-16 to represent each stimulus image (*d* = 1000), feed these features into 6-layer stacked LSTM with 1000 cells (*h* = 1000) and adopt linear transforms of 256 units (*ht* = 256) and 3 units (L, R and N, *m* = 3). The number of layers stacked in LSTM is a hyperparameter. We adopt 6-layer stacked for the reason that we expect the number of 6 could give us a good chance to match brain regions and better examine whether each LSTM layer can learn representations at different levels of abstraction.

#### Training and model evaluation

In training our deep neural network, a common cross-entropy loss function is used for optimization, estimated using the maximum likelihood method:

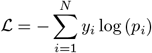

where label *y*_0_ = 1 if monkey had a leftward saccade and *y*_1_ = 1 if monkey had a rightward saccade else *y*_2_ = 1. The five stimuli on the boundary corresponded to *y*_2_ = 1 regardless of the cue shape. The Adam optimizer is used with the default learning rate of 1*e* − 5. We use a batch size of 1 (i.e., trial-by-trial learning) to match the training procedure of the monkeys. All the simulations are implemented using PyTorch [34].

To evaluate the categorization accuracy of our model, we use micro-averaged F1 score metric. The micro-averaged F1 score pools per-sample classifications across classes to compute the micro-averaged precision

(P) and recall (R) by counting the total true positives, false negatives, and false positives, and then a harmonic class of P and R, as follows

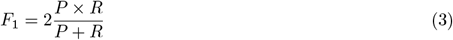

ranges between 0 and 1. The closer it is to 1, the better the model. Our model achieves the F1 score of 98% at least in multiple sampling frames (Supplemental Fig. S2).

#### Representational similarity analysis

In order to relate our artificial deep neural network to the brain, we utilize representational similarity analysis (RSA), an experimental and analytical framework, relating representations in the brain and our model by computing and comparing representational dissimilarity matrices (RDMs) that characterize the information encoded in a given brain or model. Each 16 *×* 16 RDM in this study contains distances/dissimilarities for pairwise stimuli. The distance indicates how dissimilar those two stimuli are in the monkey brain or how dissimilar those two model representations are in the model.

#### Candidate neural dissimilarity measures

There are many possible measures to calculate the distance/dissimilarity between two or more task-relevant and stimuli-relevant spike trains in neural data. In this study, we consider candidate measures, such as rate coding, inter-spike interval (ISI) distance and SPIKE distance. Rate coding considers spike count within a time bin and represents each stimulus as a neural firing rate vector which consists of spike count across neurons whereas ISI and SPIKE distance take spike timing into account. ISI emphasizes the relative intervals between spikes while SPIKE also incorporates the absolute timing of spikes which can be useful for evaluating synchrony between spike trains. Pairwise-stimuli ISI/SPIKE distance is calculated by temporal averaging over a period of trial time. Specifically, rate coding explicitly constructs representation vector for individual stimulus before constructing the RDM using Euclidean distance, cosine distance or Pearson correlation, and the other two measures construct pairwise statistics towards RDM directly. More detailed information of candidate neural dissimilarity measures, please see Supplementary Information.

When processing neural data, we extract the spike trains of isolated neurons detected under electrodes. Regardless of which candidate measure we use, each element in task-relevant RDMs is the averaged distance (dissimilarity) from spike trains (in the time bins) across task-relevant trials and monkeys using a python toolbox PySpike [35] implemented ISI-distance and SPIKE-distance for the numerical analysis of spike train dissimilarity.

#### Measure evaluation

To identify the best neural dissimilarity measure to analyze neural data, we opt for a RSA-based approach where we correlate RDMs derived from candidate neural dissimilarity measures to a static reference matrix of stimuli. This reference matrix reflects experimenter’s intended relationship between stimuli and we assume that the best measure should be the one that yields the highest Spearman’s rank correlation to the reference matrix. To construct such a reference matrix *E*, we represent each stimulus using experimenter-indented coordinates and compute pairwise dissimilarities. 16 color-motion stimuli are distributed in four quadrants of a coordinate system where each stimulus has a unique x-y coordinate. For example, bottom-left item in the coordinate system is denoted as (1, 1) and the top-right item is denoted as (4, 4) in Fig. 1c. For any two stimuli *a* and *b* with coordinates (*a*_*x*_, *a*_*y*_) and (*b*_*x*_, *b*_*y*_), we compute their distance/dissimilarity *E*(*a, b*) as

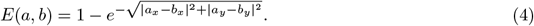

We derive such an experimenters-intended-based RDM, *E*, by calculating the exponential distance of the coordinates of pairwise color-motion stimuli. Note that being subtracted by 1 and the exponential function have no effect on the calculation of Spearman rank correlation. Although experimenter-intended-based RDM provides a fixed reference in both motion and color contexts, it inherently incorporated task-relevant distinctions in how it organized information based on stimuli coordinates.

#### RDM based on model representations

To derive task-relevant RDMs from model representations in the deep neural network, we fill the entries by taking the averaged distance (dissimilarity) between hidden states in the LSTM across task-relevant trials. Although we employ Euclidean distance to calculate dissimilarities, the results were not affected by using Cosine distance (Extended Data Fig. 5).

#### LSTM time estimated

Given that the stacked LSTM processes stimulus images in a sequential manner, our model contains hidden states that evolve dynamically over trial time (i.e., stimulus images). By capturing the hidden states from processing the first image (fixation) to the last image (moving colored stimulus), we can derive *n* task-relevant RDMs from trials, where *n* is the number of stimulus images. To relate with sliding windows, we use LSTM time estimated by the frames of images. By selecting 60 frames per second, each image is displayed for 1*/*60 of a second, allowing the input of each image to be considered as a time stamp (Extended Data Fig. 1).

#### Dimensional stretching

In two categorization tasks, dimensional stretching refers to the expansion of the perceived psychological difference beyond the physical difference between stimuli. In our study, attentional effects refer to how the monkeys allocate attention to relevant dimension in response to their saccade response in task-relevant context. We evaluate dimensional stretching through model-free and model-fit approaches to gain insight into how the monkey brain reconfigured along motion-relevant and color-relevant dimensions in task-relevant context.

#### Dimensional stretching: a model-free approach

The model-free approach evaluates dimensional stretching without assuming an underlying psychological space. Simply, we compare the dissimilarity between item pairs mismatch on one dimension and match on the other under two task contexts.

We denote the dissimilarity between item pairs (*i, j*) as *D*_*m*_(*i, j*) and *D*_*c*_(*i, j*) that mismatch on motion and mismatch on color, respectively, for instance, in Fig. 1c, stimuli in the bottom-left corner (pink and upward) and stimuli in the top-left corner (green and upward) mismatch on color and match on motion. For *N* (*i, j*) = 24 pairs under motion-relevant context (i.e., (*i, j*)^*m*^) and color-relevant context (i.e., (*i, j*)^*c*^), we perform two-tailed paired *t* -test to show the difference of pairs that in different tasks, e.g., *D*_*c*_(*i, j*)^*c*^ − *D*_*c*_(*i, j*)^*m*^ and *D*_*m*_(*i, j*)^*m*^ − *D*_*m*_(*i, j*)^*c*^.

#### Dimensional stretching: a cognitive model

The model-fit approach aims to explicitly model the psychological space itself, which assumes attentional effects exist and finds the best-fit attention for task-relevant context. We use a cognitive model that hypothesizes attention weights towards task-relevant and task-irrelevant dimension, then derive an attentional RDM, *AE*. Specifically, we assign the stimulus coordinates and for pairwise stimuli item *a* and *b*, the distance/dissimilarity in the task content, for example, the horizontal dimension is the task-relevant dimension, is calculated as

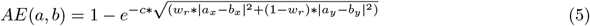

where *c* is constant, *w*_*r*_ represents the attention weight in the dimension relevant to the task and *w*_*r*_ *>*=0.We find best-fit parameters *w*_*r*_ to the task-relevant dimension by maximizing the Spearman rank correlation between attentional matrix *AE* and representation-based RDM with the grid search,

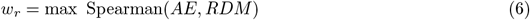

where all stimuli items are fit at once to provide one estimate of stretching along the dimension of color and motion. Based on the cognitive model, we can explore how attention weights change depending on the task cued.

#### Quantification and statistical analysis

In general, we assume that data were normally distributed but this is not formally tested. We conduct a one-way ANOVA analysis to validate the significance of the type of measure, and employ a two-tailed paired *t* -test with Bonferroni correction for pairwise comparisons to demonstrate that ISI outperforms other measures. In order to evaluate adaptive stretching, we utilize a two-tailed paired *t* -test and calculate the p-value to establish statistical significance. We perform 1000 permutation tests by shuffling task labels to assess how extreme the attention weight is in that distribution is compared to the null distribution. All statistical tests are described in the main text. Levels of statistical significance are indicated as follows: ^*^*p <* .05, ^**^*p <* .01, ^***^*p <* .001, ^****^*p <* .0001.

## Supplementary Information for

### Simulating trials

In the original experiment [1], two rhesus monkeys, one female (Paula) and one male (Rex), were trained to perform two categorization tasks. Monkeys learned to categorize stimuli into different classes indicated by the task cue. The cued task was color in half the trials and motion in the other half. Specifically, monkeys were shown a sequence of animated movies that are made of a fixation point (0.5 s), a cue (1 s), and stimulus (up to 3 s) in each trial. The cue reflects whether stimuli should be categorized by its motion or color. There are four types of cues, namely the cross, the quatrefoil, the circle and the triangle cue. The cross and the quatrefoil indicate that stimuli categorization should be made based on motion and the circle and the triangle indicate stimuli categorization should be made based on color. After cueing, the cue disappeared and the stimulus will be presented at the center of the fixation spot. Monkeys were free to respond any time up to 3 s after stimulus onset. During learning, neuronal activity was recorded in six regions of the brain across a maximum of 108 electrodes simultaneously. Images of the fixation point and cue are static in movies, while stimulus images are moving, made of 400 moving dots with one out of 21 color-motion combinations and one out of 2 dot’s speed. There are 21 color-motion stimuli from 7 possible colors and 7 possible motion directions (Fig. 1c). For correct responses to categorization, monkeys were rewarded with apple juice. There are 16 color-motion stimuli not near the category boundary but 5 ambiguous stimuli items on or near the category boundary. Monkeys were always rewarded for not giving any responses in ambiguous trials (stimuli on the category boundary). These ambiguous trials were excluded for calculating the animals’ percent correct performance in the original experiment.

To train our deep learning model, we construct identical fixation and cue images to the ones in the original experiment and stimulus images with the same stimuli patterns (stimulus diameter, dot diameter, number of dots and dot speed). Since images of the fixation point and cue are static, we repeat the same image to simulate the movies shown to monkeys, while the stimuli dots are moving, thus we choose a frequency of 60 frames per second to make it look continuous, that is a total of 270 images with 30 images of the fixation point, 60 images of cue and 180 images of stimuli in each trial. (The sampling frequency has no impact on model accuracy, please see Supplemental Figure S2).

### Stimulus image details

We construct 128*×*128 pixels RGB images. In the fixation point image, the distance between the fixation point and either of the grey dots is equal to 1*/*4 of the horizontal length of the image. This ratio is kept constant for the rest of the image types. We import the monochrome cue images and changed the cue color to grey, the background color to black, adding the three dots also appearing in the fixation point image. Stimuli are colored dynamic random dot patterns with 100% motion coherence presented centrally on the fixation spot (stimulus diameter: 3.2; dot diameter: 0.08; number of dots: 400; dot speed: 1.67 /s or 10 /s for half of the recording trials, respectively). All colors are defined in the CIE L*a*b* space and had the same luminance and saturation.

### Measures of spike trains dissimilarity

In this work, we consider candidate neural dissimilarity measures, such as rate coding, ISI distance and SPIKE distance. Rate coding utilizes a simple scheme where total spike counts alone carry information. Measures such as ISI distance and SPIKE distance focus on the temporal profiles of brain activity. ISI distance measures relative spike timing to see whether rhythms and patterns match, while SPIKE distance also incorporates the absolute timing of spikes which can be useful for evaluating synchrony between spike trains.

**Figure S1:**
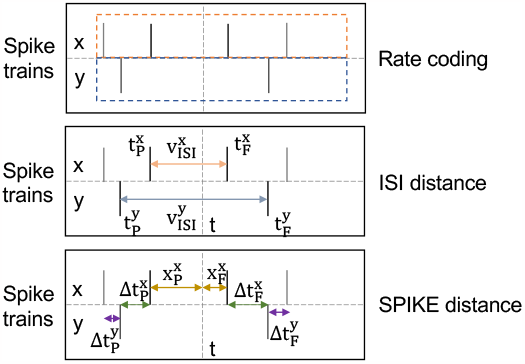
The illustration of spike trains dissimilarity measures. Rate coding (the first row) represents a stimulus as the total number of spikes across neurons. ISI distance (the second row) compares two stimuli based on the relative spike interval 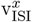 and 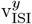 while SPIKE distance (the third row) compares two stimuli, not only based on the inter-spike interval but also on the absolute spike timing (e.g., Δt_P_, Δt_F_, x_P_ and x_F_).

**Rate coding** represents stimulus as neural firing rate across neurons in brain regions or lobes over a period of time. It is assumed that a time bin with higher firing rate represents more information. To construct rate-coding-based RDM, we first represent each stimulus as a 1D vector whose entries are the number of spikes across neurons/electrodes. We then compute pairwise distance/similarity between 16 stimuli using a range of distance/similarity measures including Euclidean, Cosine distance and Pearson correlation which fill the entries of a RDM.

**ISI distance and SPIKE distance** directly calculate the distances between stimuli pairs from spike trains. With differences which we will note below, both measures can be defined as the temporal average of the respective time profile

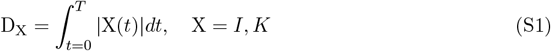

where *I* denotes ISI distance function and *K* denotes SPIKE distance function.

For each spike train *u* ∈ *{x, y}*, at each time instant (Supplemental Figure S1), the time of the previous spike is

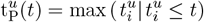

and the time of the following spike is

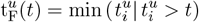

as well as the interspike interval is

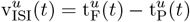

**ISI distance** focuses on the relative firing rate pattern based on the instantaneous interspike intervals [2]. The ISI distance between spike trains *x* and *y* at time instant *t* is calculated as

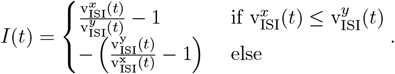

For spike trains which are identical or have constant and equal interspike intervals with phase shift, the ISI distance reaches 0. ISI distance approaches -1 or 1 if the first or the second spike train is much faster than the other. We derive ISI-distance-based RDM consisting of ISI distance between 16 stimuli by temporal averaging over the absolute values |*I*(*t*)| in the time bin using Eq. S1.

**SPIKE distance** builds upon the properties of the ISI distance with a specific focus on absolute spike timing [3], which accounts for both firing rate differences as well as precise spike time differences.

The SPIKE distance profile is calculated in two steps: first, a spike time difference is calculated for each spike and then for each time instant the relevant spike time differences are selected, weighted, and normalized. Specifically, for two spike trains *x* and *y*, each time instant is surrounded by four spikes: the preceding spike from *x* spike train 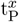, the following spike from *x* spike train 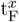, the preceding spike from *y* spike train 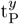 and the following spike from *y* spike train 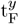 . Each of these spikes can be identified with a spike time difference to the nearest spike in the other spike train, for example, the instantaneous differences of previous and following spike times are denoted as

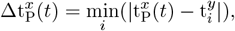

and analogously for 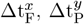 and 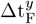 (Supplemental Figure S1). For spike train *u* ∈ *{x, y}*, the intervals from the time instant under consideration to the previous and the following spikes is

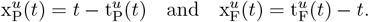

The local weighting for the spike time differences of the spike train *u* ∈ {*x, y*} is

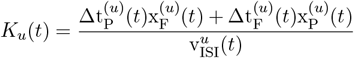

which weights the spike time differences for each spike train according to the relative distance of the corner spike from the time instant under consideration. In the last step, the two contributions of the two spike trains are locally weighted by their instantaneous interspike intervals, and then the SPIKE distance between the spike trains *x* and *y* at time instant *t* is calculated as

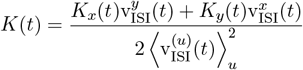

which ranges from 0 to 1. A value of 0 indicates identical spike trains and 1 indicates maximal dissimilarity. Similarly, we derive SPIKE-distance-based RDM consisting of SPIKE distance between 16 stimuli by temporal averaging over the absolute values |*K*(*t*)| in the time bin using Eq. S1.

## Supplemental Figures and Tables

**Figure S2:**
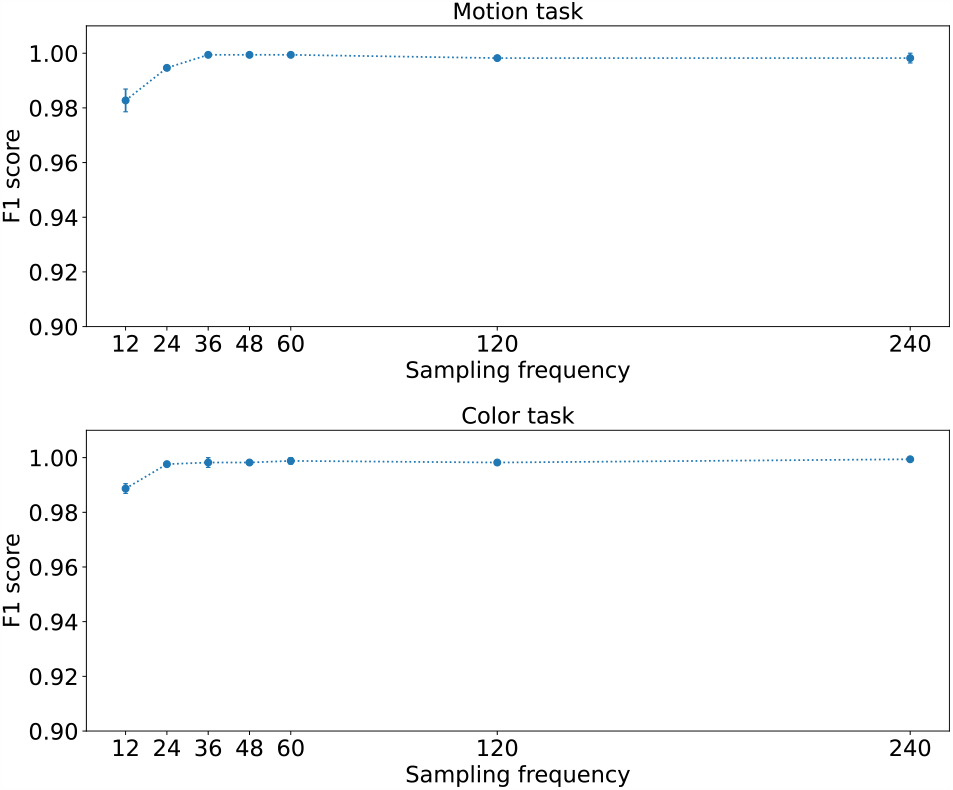
Model performance in different sampling frequency. F1 score is applied here. Each dot represents one chosen sampling frequency (i.e., frames per second) for stimulus images, and the performance score is averaged over task-relevant trails at default learning rate. The original experiment reported accuracy of 94% and 89% for the motion and color tasks in the monkey brain, respectively. In contrast, our CNN-LSTM model achieves a higher accuracy of 98% at least.

**Table S1:** The type of measure matters in region PFC (*F* (4, 105) = 81.98, *p <* .0001). Two-tailed *t* -test performed with Bonferroni correction *α* = 0.05*/n* = 0.0125 (*n* = 4) in pairwise comparison.

| $t$-test | ISI |
| --- | --- |
| SPIKE | $t(21) = 4.25; p = .00012$ |
| Euclidean(roi) | $t(21) = 12.89; p < .0001$ |
| Cosine(roi) | $t(21) = 11.83; p < .0001$ |
| Pearson(roi) | $t(21) = 11.83; p < .0001$ |

**Table S2:** The type of measure matters in region FEF (*F* (4, 105) = 260.1, *p <* .0001). Two-tailed *t* -test performed with Bonferroni correction *α* = 0.05*/n* = 0.0125 (*n* = 4) in pairwise comparison.

| $t$-test | ISI |
| --- | --- |
| SPIKE | $t(21) = 3.53; p = .0010$ |
| Euclidean(roi) | $t(21) = 26.49; p < .0001$ |
| Cosine(roi) | $t(21) = 21.84; p < .0001$ |
| Pearson(roi) | $t(21) = 21.84; p < .0001$ |

**Table S3:** The type of measure matters in region LIP (*F* (4, 105) = 29.98, *p <* .0001). Two-tailed *t* -test performed with Bonferroni correction *α* = 0.05*/n* = 0.0125 (*n* = 4) in pairwise comparison.

| $t$-test | ISI |
| --- | --- |
| SPIKE | $t(21) = 6.45; p < .0001$ |
| Euclidean(roi) | $t(21) = 5.77; p < .0001$ |
| Cosine(roi) | $t(21) = 9.16; p < .0001$ |
| Pearson(roi) | $t(21) = 8.73; p < .0001$ |

**Table S4:** The type of measure matters in region IT (*F* (4, 105) = 31.69, *p <* .0001). Two-tailed *t* -test performed with Bonferroni correction *α* = 0.05*/n* = 0.0125 (*n* = 4) in pairwise comparison.

| $t$-test | ISI |
| --- | --- |
| SPIKE | $t(21) = 14.09; p < .0001$ |
| Euclidean(roi) | $t(21) = -1.09; p = .2804$ |
| Cosine(roi) | $t(21) = 6.12; p < .0001$ |
| Pearson(roi) | $t(21) = 5.28; p < .0001$ |

**Table S5:** The type of measure matters in region MT (*F* (4, 105) = 84.66, *p <* .0001). Two-tailed *t* -test performed with Bonferroni correction *α* = 0.05*/n* = 0.0125 (*n* = 4) in pairwise comparison.

| $t$-test | ISI |
| --- | --- |
| SPIKE | $t(21) = -0.54; p=.5947$ |
| Euclidean(roi) | $t(21) = 8.27; p<.0001$ |
| Cosine(roi) | $t(21) = 13.65; p<.0001$ |
| Pearson(roi) | $t(21) = 13.19; p<.0001$ |

**Table S6:**
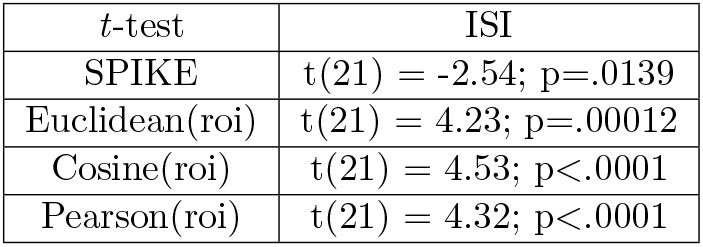
The type of measure matters in region V4 (*F* (4, 105) = 21.62, *p <* .0001). Two-tailed *t* -test performed with Bonferroni correction *α* = 0.05*/n* = 0.0125 (*n* = 4) in pairwise comparison.

| $t$-test | ISI |
| --- | --- |
| SPIKE | $t(21) = -2.54; p=.0139$ |
| Euclidean(roi) | $t(21) = 4.23; p=.00012$ |
| Cosine(roi) | $t(21) = 4.53; p<.0001$ |
| Pearson(roi) | $t(21) = 4.32; p<.0001$ |

**Table S7:** The type of measure matters in frontal lobe (*F* (4, 105) = 225.5, *p <* .0001). Two-tailed *t* -test performed with Bonferroni correction *α* = 0.05*/n* = 0.0125 (*n* = 4) in pairwise comparison.

| $t$-test | ISI |
| --- | --- |
| SPIKE | $t(21) = 5.27; p<.0001$ |
| Euclidean(lobe) | $t(21) = 21.46; p<.0001$ |
| Cosine(lobe) | $t(21) = 19.17; p<.0001$ |
| Pearson(lobe) | $t(21) = 19.66; p<.0001$ |

**Table S8:** The type of measure matters in parietal lobe (*F* (4, 105) = 16.08, *p <* .0001). Two-tailed *t* -test performed with Bonferroni correction *α* = 0.05*/n* = 0.0125 (*n* = 4) in pairwise comparison.

| $t$-test | ISI |
| --- | --- |
| SPIKE | $t(21) = 3.62; p=.0008$ |
| Euclidean(lobe) | $t(21) = 5.84; p<.0001$ |
| Cosine(lobe) | $t(21) = 6.12; p<.0001$ |
| Pearson(lobe) | $t(21) = 5.44; p<.0001$ |

**Table S9:** The type of measure matters in temporal lobe (*F* (4, 105) = 20.58, *p <* .0001). Two-tailed *t* -test performed with Bonferroni correction *α* = 0.05*/n* = 0.0125 (*n* = 4) in pairwise comparison.

| $t$-test | ISI |
| --- | --- |
| SPIKE | $t(21) = -1.14; p=.2609$ |
| Euclidean(lobe) | $t(21) = 5.93; p<.0001$ |
| Cosine(lobe) | $t(21) = 6.17; p<.0001$ |
| Pearson(lobe) | $t(21) = 5.09; p<.0001$ |

---

Model (Extended Data Figs. 1)

Result I extended data (Extended Data Figs. 2, 3, 4)

Stats for Result I (Extended Data Tables 1, 2)

Result II extended data (Extended Data Fig. 5)

Stats for Result III (Extended Data Tables 3, 4)

**Extended Data Fig. 1.**
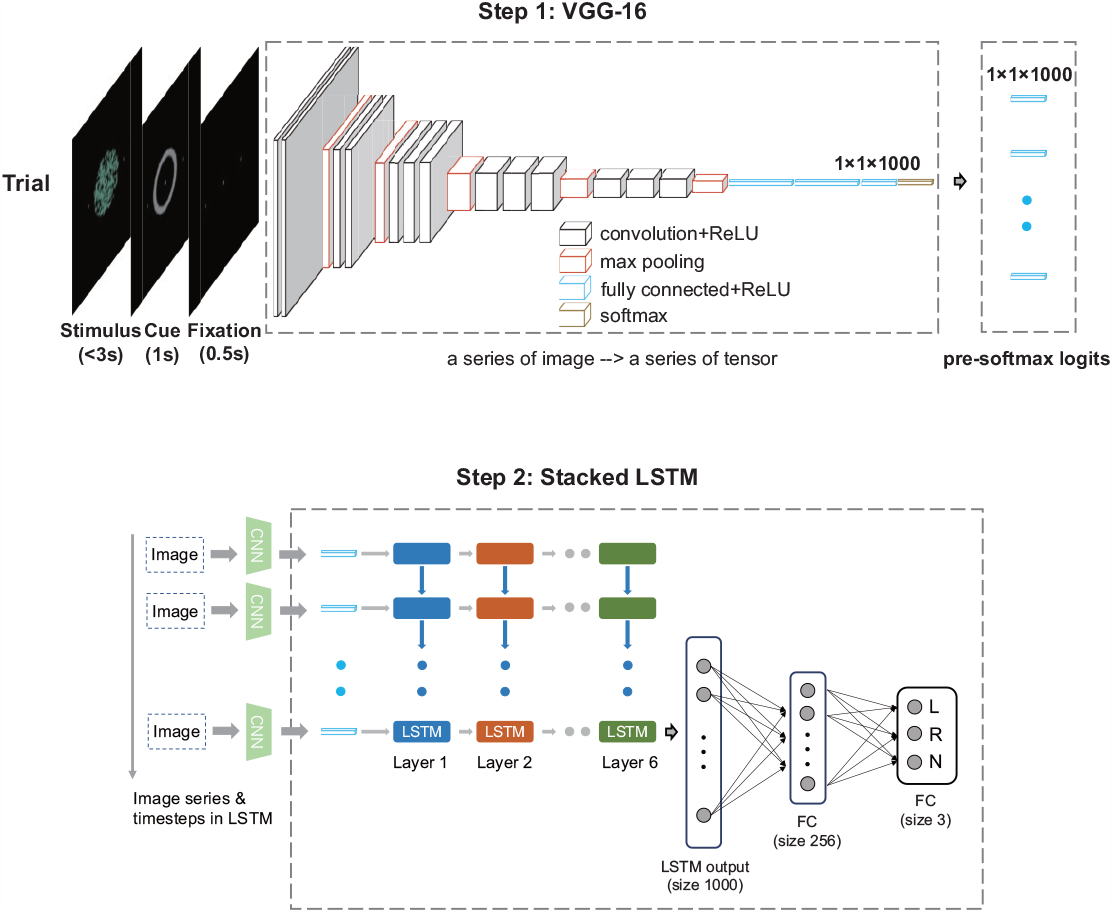
CNN-LSTM model. In our artificial neural network (ANN), a pretrained VGG-16 that maps each two-dimensional image (input) to a pre-softmax tensor of size 1000 (output). The output of the CNN, concatenated with pre-softmax logits, serves as an input to a six-layer stacked LSTM. To transform LSTM representations into the decision response, we apply two fully connected (FC) linear transformations, one with 256 units and the other with 3 units (L, R and no response). Each stimulus image is a time stamp, denoted as LSTM time.

**Extended Data Fig. 2.**
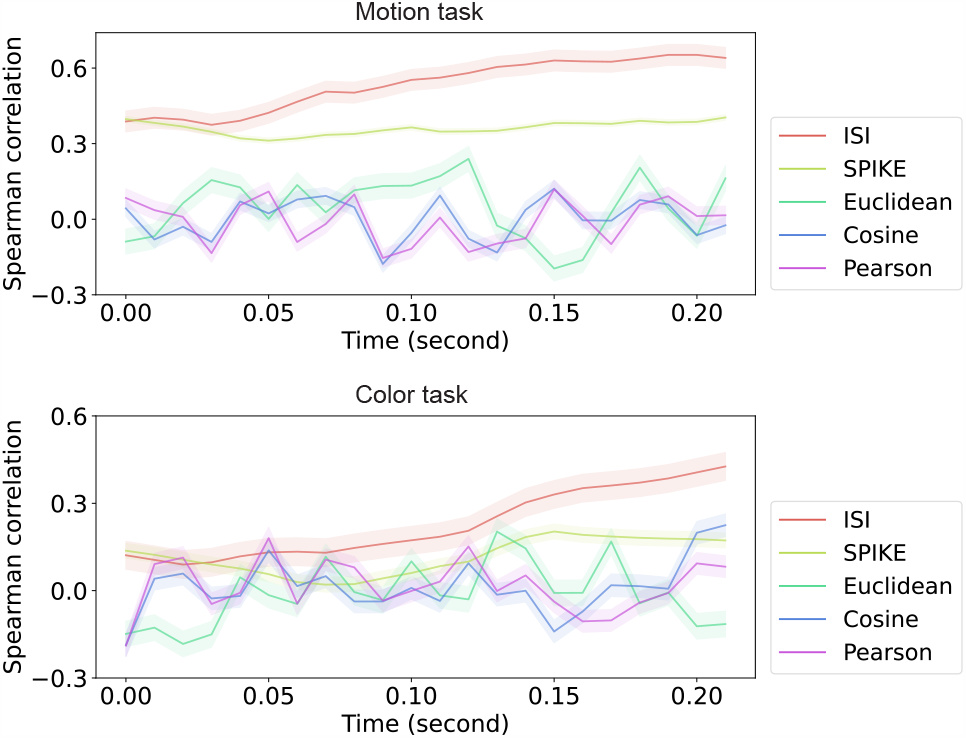
Spike timing measures best capture the experimenter intended coordinates. We compared the Spearman rank correlation between dissimilarity matrices (i.e., RDMs) constructed from the monkey data for the 16 unambiguous items with an RDM derived from the experimenter-intended stimulus coordinates. A 20 ms sliding window is used and moved in 10 ms steps. A one-way ANOVA analysis is conducted to show the significance of different types of measures, *F* (4, 105) = 124.67, *p <* .0001. ISI measure surpasses other measures and best captures the experimenter-intended coordinates. A 50 ms time window is shown in the main text, indicating the length of time window chosen does not affect the advantage of timing measures in assessing spike trains in such categorization tasks. Time on the horizontal axis is measured from stimulus onset.

**Extended Data Fig. 3.**
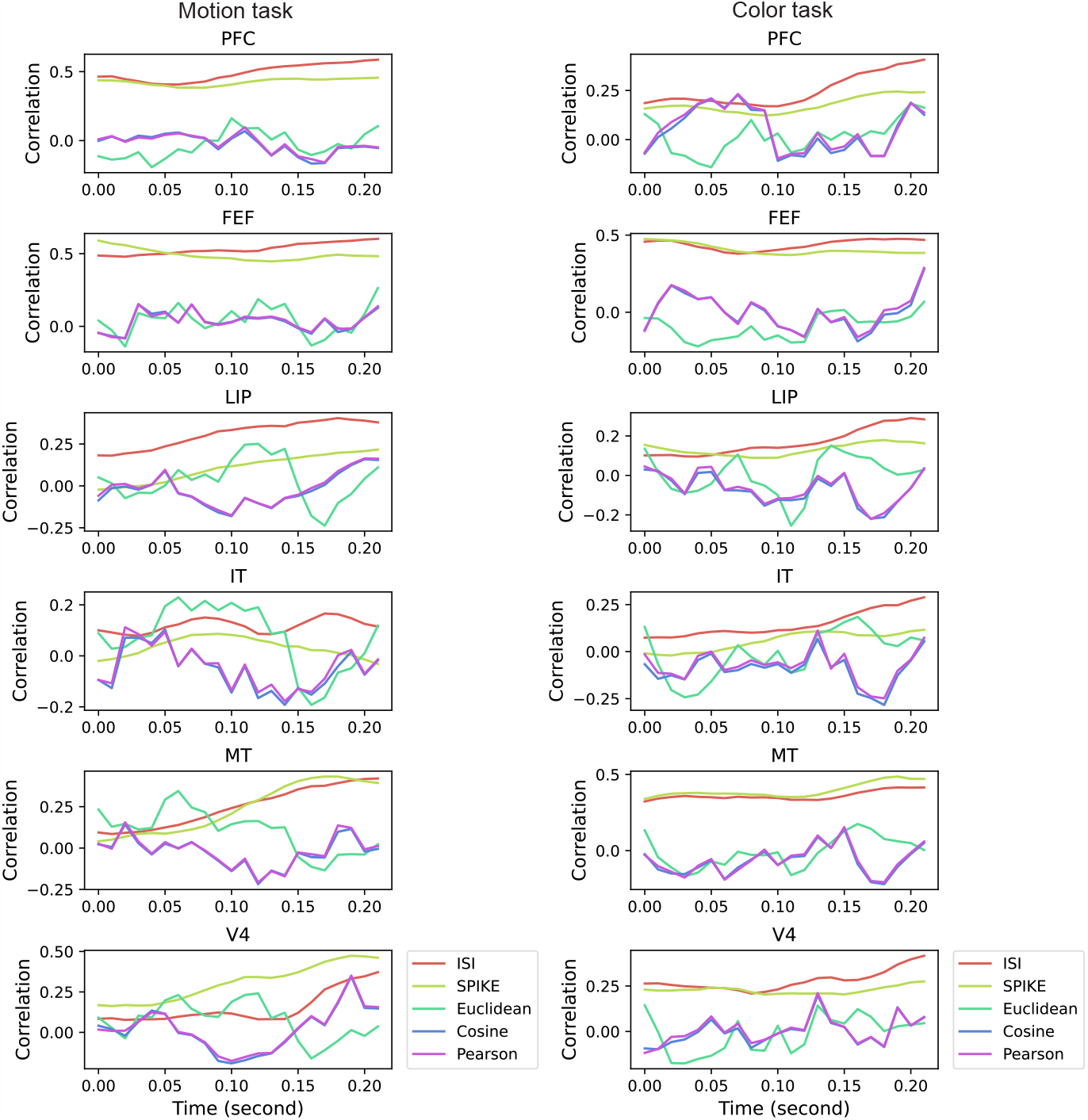
Spike timing matters in each individual brain region. We compared the Spearman rank correlation between dissimilarity matrices (i.e., RDMs) constructed from the monkey data in each individual brain region for the 16 unambiguous items with an RDM derived from the experimenter-intended stimulus coordinates. A 50 ms sliding window is used here. Spike timing measures best capture the experimenter intended coordinates in six brain regions

**Extended Data Fig. 4.**
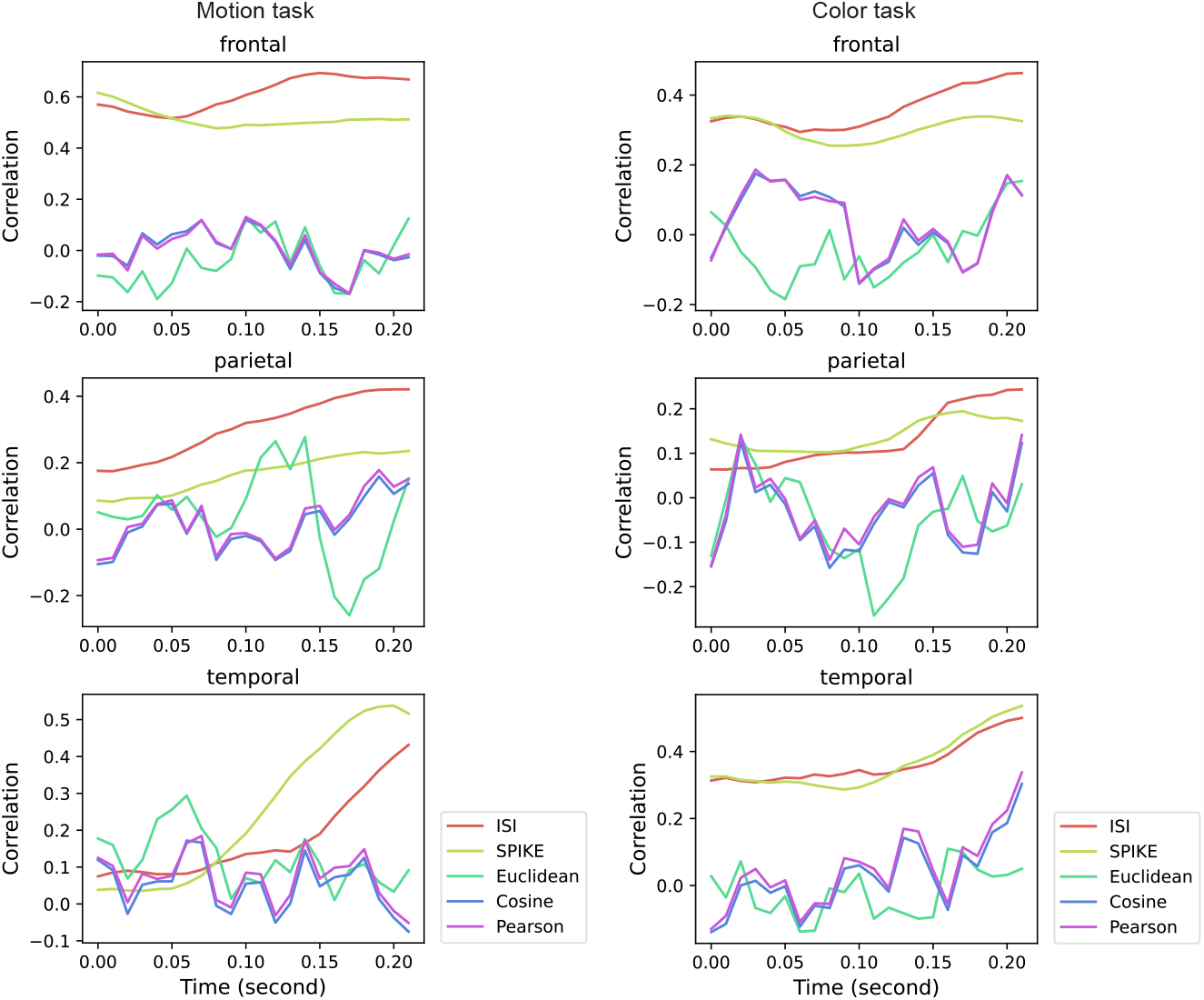
Spike timing matters in each individual lobe. We compared the Spearman rank correlation between dissimilarity matrices (i.e., RDMs) constructed from the monkey data in each individual lobe for the 16 unambiguous items with an RDM derived from the experimenter-intended stimulus coordinates. 50 ms sliding window is chosen here. Spike timing measures best capture the experimenter intended coordinates in three lobes (frontal, parietal and temporal).

**Extended Data Fig. 5.**
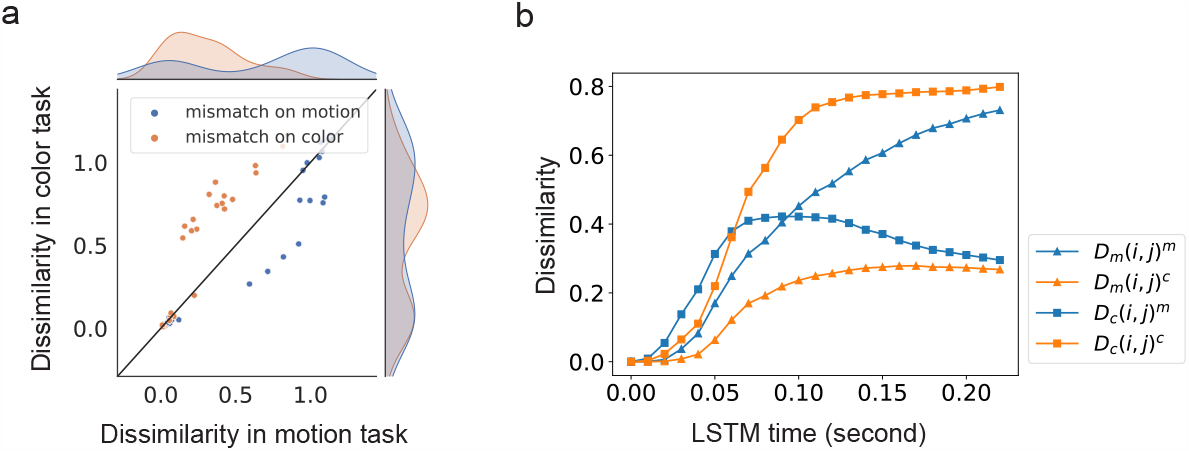
Dimensional stretching occurs in model representations with cosine distance. **a**, Dissimilarity between item pairs mismatching on motion or color is increased when that dimension is task relevant. The density distributions of these dissimilarities also indicate this task-modulation. **b**, Changes in dissimilarity over LSTM time with 60 frames per second in model representations. Stimuli pairs that mismatch on task-relevant dimension become more different in task-relevant context than task-irrelevant context, and dimensional stretching was significantly different than 0 (*D*_*c*_(*i, j*)^*c*^ − *D*_*c*_(*i, j*)^*m*^: *M* = 0.4909, *t*(23) = −8.541, *p <* .0001; *D*_*m*_(*i, j*)^*m*^ − *D*_*m*_(*i, j*)^*c*^: *M* = 0.4481, *t*(23) = −9.178, *p <* .0001).

**Extended Data Table 1.**
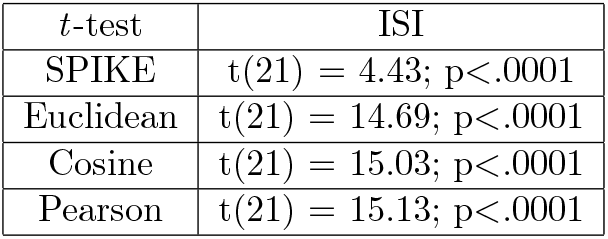
Two-tailed *t* -test with Bonferroni correction *α* = 0.05*/n* = 0.0125 (*n* = 4) was conducted to show that the ISI measure performed best (50 ms) in pairwise comparisons.

**Extended Data Table 2.** Two-tailed *t* -test with Bonferroni correction *α* = 0.05*/n* = 0.0125 (*n* = 4) was conducted to show that the ISI measure performed best (20 ms) in pairwise comparisons.

| $t$-test | ISI |
| --- | --- |
| SPIKE | $t(21) = 5.80; p < .0001$ |
| Euclidean | $t(21) = 13.00; p < .0001$ |
| Cosine | $t(21) = 14.72; p < .0001$ |
| Pearson | $t(21) = 14.16; p < .0001$ |

**Extended Data Table 3.** Permutation test statistics on attentional effects in six brain regions. We performed 1000 times to assess how extreme it is according to the null distributions built up by shuffling task labels.

| Region | Permutation test (motion task) | Permutation test (color task) |
| --- | --- | --- |
| PFC | rank=1 , p=.0020 | rank=1 , p=.0020 |
| FEF | rank=2 , p=.0040 | rank=1 , p=.0020 |
| LIP | rank=2, p=.0040 | rank=1, p=.0020 |
| IT | rank=1, p=.0020 | rank=2, p=.0040 |
| MT | rank=1, p=.0020 | rank=1001, p=.0020 |
| V4 | rank=1000, p=.0040 | rank=2, p=.0040 |

**Extended Data Table 4.** Permutation test on attentional effects in LSTM multilayers. We performed 1000 times to assess how extreme it is according to the null distributions built up by permuting task labels.

| Layer | Permutation test |
| --- | --- |
| 1 | rank=164, p=.3277 |
| 2 | rank=8, p=.0160 |
| 3 | rank=5, p=.0100 |
| 4 | rank=3, p=.0060 |
| 5 | rank=1, p=.0020 |
| 6 | rank=1, p=.0020 |

